# Hippocampal blood flow rapidly and preferentially increases after a bout of moderate-intensity exercise in older adults with poor cerebrovascular health

**DOI:** 10.1101/2022.07.15.500132

**Authors:** Jacqueline A. Palmer, Jill K Morris, Sandra A. Billinger, Rebecca J Lepping, Laura Martin, Zachary Green, Eric D. Vidoni

**Affiliations:** Department of Neurology, School of Medicine, University of Kansas Medical Center, Kansas City, KS, United States of America; University of Kansas Alzheimer’s Disease Research Center, Fairway, KS, United States of America; Department of Molecular & Integrative Physiology, University of Kansas Medical Center, Kansas City, KS, USA; Department of Physical Therapy, Rehabilitation Science, and Athletic Training, School of Health Professions, University of Kansas Medical Center, Kansas City, KS, United States of America

**Keywords:** cardiovascular, cerebral blood flow, arterial spin labelling MRI, neurovascular, vascular plasticity

## Abstract

Over the course of aging, there is an early degradation of cerebrovascular health that may be attenuated with aerobic exercise training. Yet, the acute cerebrovascular response to a single bout of exercise remains elusive, particularly within key brain regions most affected by age-related disease processes. We investigated the acute global and region-specific cerebral blood flow (CBF) response to 15 minutes of moderate-intensity aerobic exercise in older adults (≥65years) (n=60) using arterial spin labeling magnetic resonance imaging. Within 0-6 minutes post-exercise, CBF decreased across all regions, an effect that was attenuated in the hippocampus. The exercise-induced CBF drop was followed by a rebound effect over the 24-minute post-exercise assessment period, an effect that was most robust in the hippocampus. Individuals with low baseline perfusion demonstrated the greatest hippocampal-specific CBF effect post-exercise, showing no immediate drop and a rapid increase in CBF that exceeded baseline levels within 6-12 minutes post-exercise. Gains in domain-specific cognitive performance post-exercise were not associated with changes in regional CBF, suggesting dissociable effects of exercise on acute neural and vascular plasticity. Together, the present findings support a precision-medicine framework for the use of exercise to target brain health that carefully considers age-related changes in the cerebrovascular system.

## 1. Introduction

Repeated engagement in aerobic exercise training stimulates potent neuroprotective effects and robust vascular plasticity that benefit neural function and cognitive behavior over the course of aging (Ahlskog et al., 2011; Cotman & Berchtold, 2002; Vecchio et al., 2018; Zhang et al., 2020). Older adults with poor vascular health and at high-risk for disease appear to show the most robust cognitive benefits from chronic exercise interventions (Ekelund et al., 2020; Ngandu et al., 2015). In particular, long-term aerobic exercise elicits potent neural and vascular plasticity in hippocampal brain structures (Erickson et al., 2011; Maass et al., 2015; Pereira et al., 2007; Seoane et al., 2022; van Praag et al., 2005), counteracting age-related loss in volume and function of hippocampal regions observed in the preclinical stages of aging disease processes (Jack et al., 2010; Raz et al., 2005). The behavioral benefits of aerobic exercise training likely occur, in part, through increases in cerebral blood flow (CBF) (Pereira et al., 2007), which declines over the course of normal aging (Alwatban et al., 2021; Xing et al., 2017). Yet, acute dynamics of immediate CBF adaptations induced by a single bout of aerobic exercise have not been well characterized. It remains unknown whether acute exercise-induced CBF mechanisms interact with aging neurobiological processes, limiting clinical translation. Such understanding is imperative for the development of innovative behavioral approaches that maximize the neuroprotective effects of exercise for aging brain health to preserve neural function and cognition.

Over the course of aging, there is a decline in CBF that occurs as early as middle age (Xing et al., 2017) and may be attenuated by engagement in regular physical activity (Sugawara et al., 2020; Thomas et al., 2013). Clinically, low resting cerebral perfusion levels in older individuals is an indicator of poor cerebrovascular health (Nagai et al., 2010; Weijs et al., 2022) and a prognostic biomarker for the development of cognitive impairment and dementia in the preclinical stages of disease (Weijs et al., 2022; Wolters et al., 2017). Lower cerebral perfusion and dysfunctional CBF regulation has been linked to age-related neural dysfunction (Stefanidis et al., 2020), cognitive impairment (Weijs et al., 2022; Wolters et al., 2017), and neurodegenerative diseases such as Alzheimer’s disease (Iadecola, 2004; Ouellette & Lacoste, 2021; Xie et al., 2016). Despite the striking effect of long-term aerobic exercise to promote cerebrovascular health, neural plasticity, and cognitive function (Zhang et al 2020; Vecchio 2018; Ahlskog 2011; Angervaren 2008), little is known about the acute onset of the first vascular and neural plastic adaptations following each bout of aerobic exercise that accrues over the course of long-term exercise interventions. Such knowledge would inform our understanding of the salient exercise-induced physiologic adaptations that may trigger beneficial changes in brain health and function over time, potentially enhancing the development of effective exercise interventions for aging populations.

Hippocampal brain structure and cognitive functions involving regional hippocampal networks appear to preferentially benefit from the long-term effects of exercise training (Erickson et al., 2011; Maass et al., 2015; Pereira et al., 2007; Seoane et al., 2022; van Praag et al., 2005). The hippocampus is one of the first brain structures to be affected by age-related disease processes (Frisoni et al., 2010; Mueller et al., 2010), with hippocampal atrophy and dysfunction beginning in late adulthood (Erickson et al., 2012; Jack et al., 1998). Age-related hippocampal changes have been linked to a decline in regional vascular health (Dhikav & Anand, 2011). These hippocampal changes in the preclinical stages of age-related disease processes are predictive for the development of mild cognitive impairment and dementia at an earlier age (Dhikav & Anand, 2011, 2007; Erickson et al., 2010; Jack et al., 1998). Long-term habitual exercise and associated higher levels of physical fitness may counteract age-related reductions in hippocampal volume and are associated with higher cognitive memory function (Erickson et al., 2009, 2011). Notably, the chronic exercise-induced increase in regional hippocampal perfusion in the aging brain appear to interact with an individuals’ baseline cardiovascular health and genetic risk for Alzheimer’s and cardiovascular disease, with long-term exercise interventions potentially benefitting those at higher genetic risk with poor baseline health to a greater degree than their lower-risk, healthy, and age-matched counterparts (Kaufman et al., 2021). While it is becoming increasingly clear that physiologic interactions between aging, baseline health, and individual genotype influence the effectiveness of exercise interventions for brain function over a long-term time scales, it remains unknown whether such interactions are detectible in the acute response to a single exercise bout. These acute physiologic interactions could potentially serve as a predictive biomarker to identify individuals with the greatest potential benefit of exercise-induced effects on brain health, particularly in vulnerable aging brain regions such as the hippocampus.

A single bout of aerobic exercise can transiently sharpen cognitive performance acutely post-exercise (Lebeau et al., 2022), though the underpinning exercise-induced physiologic mechanisms for short-term cognitive benefit remain elusive. Accumulating evidence suggests that the subtle and transient cognitive effects immediately following cessation of an exercise bout may be particularly beneficial with aging (Barella et al., 2010; Hyodo et al., 2012; Johnson et al., 2016) and across cognitive domains of episodic memory and processing speed (Davranche et al., 2009; Kamijo et al., 2007, 2009; Kao et al., 2017; Peiffer et al., 2007). Long-term improvements in cognitive function with chronic and habitual exercise are associated with increased hippocampal volume and perfusion (Jonasson et al., 2016; Maass et al., 2015; Makizako et al., 2015; Pereira et al., 2007; Thomas et al., 2013). However, it remains unknown whether short-term exercise-induced cognitive performance gains may occur through similar vascular mechanisms of increased hippocampal blood flow.

In this study we sought to characterize the acute global and region-specific CBF dynamics in response to a single bout of moderate-intensity aerobic exercise in older adults. We tested the potential interactive effects of baseline cerebrovascular health on exercise-induced regional CBF changes and explored associations with cognitive performance acutely post-exercise. We measured regional CBF at baseline and immediately after a 15-minute bout of moderate-intensity aerobic exercise (45-55% age-predicted heart rate reserve) for 24 minutes using MR-based pseudo-continuous arterial spin labeling (pCASL) (White et al., 2021). We hypothesized that hippocampal brain regions would show the most robust increases in CBF in response to exercise. Further, we predicted that older individuals with the most age-related decline in regional cerebrovascular perfusion at baseline would show the greatest exercise-induced increases in hippocampal blood flow. Finally, we hypothesized that there would be a positive relationship between change in regional-specific CBF and change in domain-specific cognitive performance acutely following exercise.

## 2. Materials and Methods

The study design is detailed in **Figure 1**. A detailed trial protocol has been described in White et al (2021) and primary endpoint analyses have been described previously (Vidoni et al., 2022).

**Figure 1.**
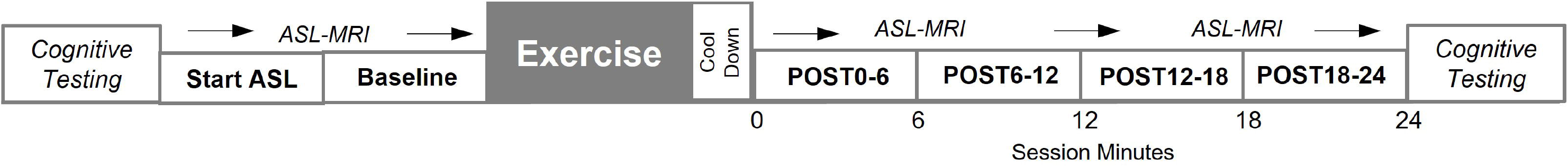
Study paradigm illustrating time course of study session, including cognitive testing and arterial spin labeling MRI measures recorded at baseline and immediately after a 15-minute bout of moderate-intensity exercise.

### 2.1. Participants

Participants (65-85 years old) were cognitively normal and possessed no significant musculoskeletal pain with exercise, MRI contraindications, psychiatric or neurological disorder, or myocardial infarction and/or symptomatic coronary artery disease in the prior 2 years. The experimental protocol was approved by the University of Kansas Medical Center Institutional Review Board (IRB#: STUDY142822) and experiments were undertaken with the understanding and written consent of each participant. Participants were genotyped for Apolipoprotein E4 (APOE4) carriage using standard procedures (White et al., 2021). The study was registered in ClinicalTrials.gov (NCT04009629).

### 2.2. General Protocol Flow

Participants attended a single visit at the MRI facility. The experimental protocol has been described in detail previously (White et al., 2021) (**Figure 1**). Briefly, in a quiet room, participants were first administered a standard cognitive battery lasting approximately 20 minutes. Then, participants changed into MRI compatible clothing and were escorted to the imaging suite. A beat-to-beat blood pressure monitor was fitted to their left index finger and calibrated. They underwent approximately 2 minutes of MR image collection. They were then escorted to an exercise cycle ergometer in an adjacent suite for 15 minutes of moderate intensity exercise. Following the exercise bout, and a 5-minute cooldown period, participants were immediately escorted back to the MR suite, and post-exercise imaging, lasting approximately 24 minutes was performed. Finally, participants returned to the cognitive testing room for the post-exercise cognitive test consistent of the same battery as at the beginning of the visit. Additional details on each step are presented in the following sections.

### 2.3. Single bout of moderate-intensity aerobic exercise on a cycle ergometer

To initiate the exercise, participants first rested on the upright ergometer for 5 minutes where resting blood pressure and heart rate were assessed. Participants then began a 5-minute warm-up followed by a 15-minute acute bout of moderate-intensity aerobic exercise on a cycle ergometer. The target intensity of 45-55% of heart rate reserve was based on age-predicted heart rate maximum (White et al., 2021) and the resistance was continually titrated to keep heart rate within the target range. A 5-minute cooldown bout at a self-selected pace with no resistance followed the exercise bout.

### 2.4. Neuroimaging procedures

MR imaging was collected before and after the exercise bout. Before exercise, two pCASL sequences (Kilroy et al., 2014; Wang et al., 2015; (Yan et al., 2010) were acquired (total scan time for each segment = 5:48, 2 M0 images). Then, a T1-weighted, 3D magnetization prepared rapid gradient echo (MPRAGE) structural scan was collected (TR/TE = 2300/2.95 ms, inversion time (TI) = 900 149 ms, flip angle = 9 deg, FOV = 253 × 270 mm, matrix = 240 × 256 voxels, voxel in-plane resolution = 1.05 × 1.05 150 mm2, slice thickness = 1.2 mm, 176 sagittal slices, in-plane acceleration factor = 2, acquisition time = 5:09).

Immediately following the exercise bout, participants returned to the scanner and completed 4 consecutive pCASL sequences, yielding a total of approximately 24 minutes of consecutive post-exercise cerebral blood flow data. All pCASL sequences were collected with the same background suppressed 3D GRASE protocol (TE/TR = 22.4/4300ms, FOV = 300 × 300 × 120mm3, matrix = 96 × 66 × 48, Post-labeling delay = 2s, 4-segmented acquisition without partial Fourier transform reconstruction, readout duration = 23.1ms, total scan time 348s, 2 M0 images).

To estimate CBF we used an estimation pipeline adapted from the Laboratory of Functional MRI (loft-lab.org, ver. February 2019). Briefly, motion artifact in labelled and control pCASL images was corrected separately. Then, average estimated CBF in each pCASL sequence was registered to the anatomical image and smoothed using 6mm full-width half maximum Gaussian kernel.

We used a regional-specific brain structure approach in our present analysis, which is in line with recent meta-analyses demonstrating age-related decline in global CBF is driven by region-specific brain structures (Weijs et al., 2022). Additionally, region-specific cerebral blood flow has been linked to domain-specific cognitive function in aging individuals (Weijs et al., 2022). Whole-cortical gray matter and 3 regional-specific brain segments (primary motor cortex, superior parietal cortex) were identified a priori based on their distinguishable functional involvement in cognitive performance domains shown to acutely benefit from exercise, notably episodic memory, selective attention, and cognitive processing speed (Davranche et al., 2009; Kamijo et al., 2007, 2009; Kao et al., 2017; Peiffer et al., 2007). To adjust for individual CBF differences and allow exploration of the temporal pattern of blood flow independent of absolute CBF, we computed regional post-exercise gray matter CBF as a percentage of baseline. We first performed individual segmentation of each anatomical scan using the Statistical Parametric Mapping CAT12 (neuro.uni-jena.de/cat, r1059 2016-10-28) package (Dahnke et al., 2013). We then isolated regions of interest as the overlap between regional gray matter segmentations and regions of interest defined by the Neuromorphometric atlas in the CAT12 toolbox {https://www.biorxiv.org/content/10.1101/2022.06.11.495736v1.full}. Average CBF in each region of interest was extracted for each pCASL sequence (github.com/aimfeld/Neurotools). Post-exercise CBF was divided by the second pre-exercise sequence CBF.

### 2.5. Cognitive behavior assessment

We employed cognitive testing for general cognitive processing speed and episodic memory on an electronic tablet using the NIH Toolbox (Kramer et al., 2014; *NIH Toolbox*, n.d.; Weintraub et al., 2013) at baseline and ~ 24 minutes post-exercise **(Figure 1**). We computed the fully corrected T-scores provided by the Toolbox, which are adjusted for formal educational attainment, age, and self-identified race and ethnicity. We used the Pattern Comparison Processing Speed Test to index general cognitive visuomotor processing speed, primarily subserved by superior parietal and primary motor cortical brain regions (Goldenkoff et al., 2021; Hatsopoulos & Suminski, 2011; Kalaska, 1996; Mutha et al., 2011; Sabes, 2011). We used the Picture Sequence Memory Test to capture episodic memory performance, with more targeted recruitment of hippocampal brain regions (Das et al., 2019; Moscovitch et al., 2016).

### 2.6 Statistical analyses

To assess whole cortical and regional gray matter CBF change over time we tested linear mixed effects models with a random intercept coefficient for each participant. P-values were obtained by likelihood ratio tests of the full model against the model without the interaction or factor in question. Age, gender, *APOE4* carrier status and mean arterial blood pressure were entered into the models. These analyses were performed using R (base and lme4 packages) (R Core Team. Vienna Austria v4.1.3 including base and lme4 packages). Relationships between change in regional CBF and domain-specific cognitive performance were explored with Pearson correlation coefficients. We used an a priori level of significance of 0.05.

## RESULTS

A total of 60 participants were included in present analyses (65% female, 72.8 ± 5.2yo) (**Table 1**).

**Table 1.** Participant characteristics.

| Table 1. Participant characteristics |  |
| --- | --- |
|  | ALL (n=60) |
| Age (years) | 72.8 ± 5.2 |
| Female, n | 39 (65%) |
| Total Workload (kilojoules) | 627±300 |
| *APOE4* carrier, n | 21 (35%) |
| Baseline Memory (fully corrected T-score) | 52.7 ± 9.9 |
| Baseline Processing Speed (fully corrected T-score) | 49.0 ± 14.1 |

Values are depicted as mean ± SD unless otherwise noted; [Range]

### 3.1. Acute exercise effects on global and regional cerebral blood flow

There was a main effect of time on CBF in the cortical gray matter (X^2^ = 81.4, p<0.001) with blood flow sharply decreasing immediately following exercise cessation followed by a steady increase over time after exercise. There was a time-by-region interaction (X^2^ = 6.1, p=0.048, **Figure 2**), in which the hippocampus showed a differential CBF response acutely after exercise compared to other brain regions. Immediately following exercise cessation (0-6 minutes post-exercise), the hippocampus showed an attenuated drop that was followed by a rapid and robust post-exercise CBF rebound effect, with CBF exceeding baseline levels by 18-24 minutes post-exercise (p=0.037). Superior parietal and motor cortical regions showed a greater drop in CBF immediately after exercise (p<.001), that was followed by a CBF rebound effect that never exceeded baseline levels at any post-exercise time point (p>0.10).

**Figure 2.**
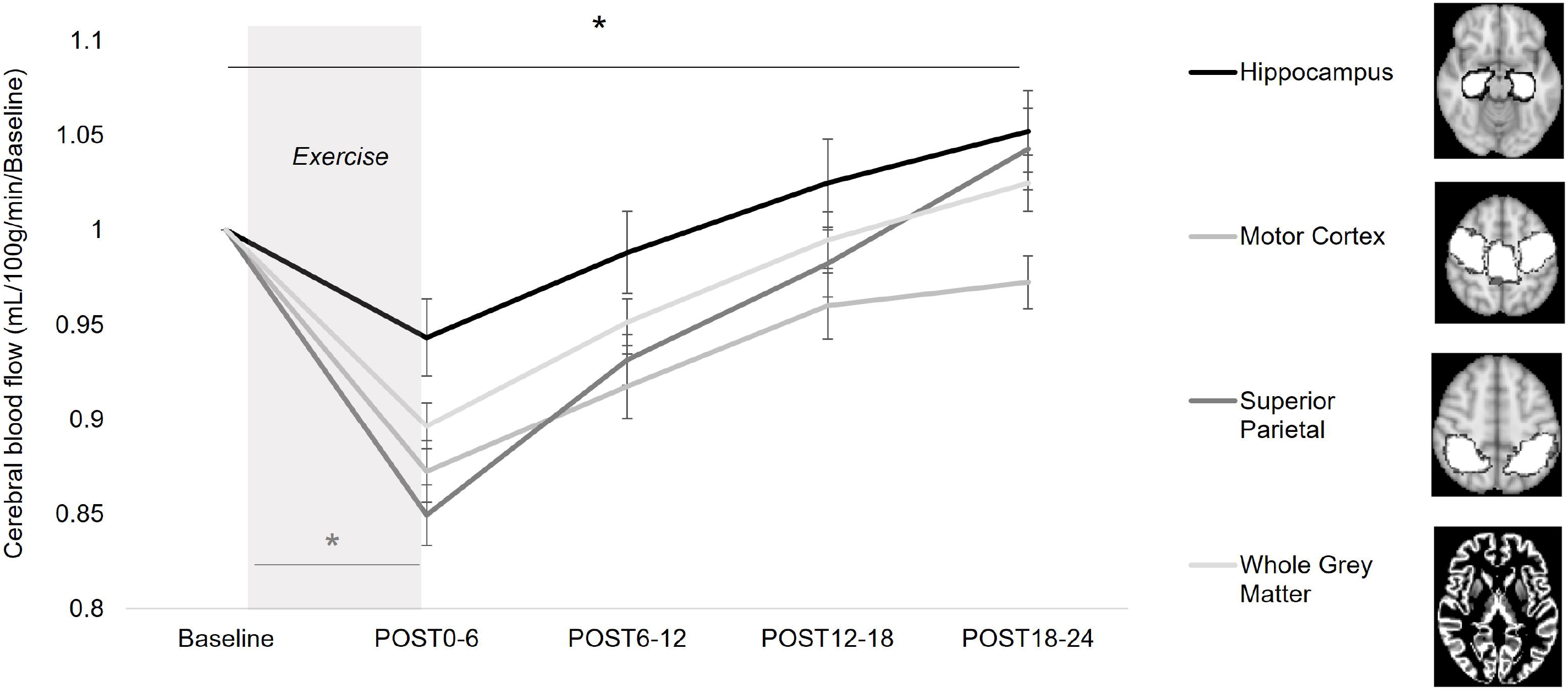
Regional cerebral blood flow (CBF) at baseline and over an acute time course following a bout of moderate-intensity aerobic exercise in older adults. All brain regions showed change in CBF over time after exercise (p<0.001), characterized by an immediate CBF reduction followed by a gradual CBF increase over the course of the 24-minute assessment period. There was a time-by-region interaction effect in which the hippocampus demonstrated an attenuated immediate CBF reduction and more robust rebound effect compared to other brain regions (X2 = 6.1, p=0.048). At the final assessment 18-24 minutes post-exercise (POST18-24), only hippocampal CBF exceeded pre-exercise baseline levels (*p=.037).

### 3.2. Differential hippocampal blood flow response to exercise as a function of baseline cerebrovascular health

To further investigate this hippocampal hyperemic response and the effect of regional cerebrovascular health in older adults, we dichotomized the cohort around median hippocampal perfusion at baseline (26.00mL/100g tissue/min). We observed a time-by-baseline perfusion (low, high) interaction of hippocampal CBF response to exercise (X^2^ = 21.1, p<0.001, **Figure 3**), in which hippocampal blood flow responded differently over time between individuals who possessed low versus high regional baseline perfusion. Post-hoc analyses revealed the region-specific hippocampal response to exercise (**Figure 2**) was driven by individuals with low baseline perfusion; here, individuals with low baseline perfusion showed an attenuated drop compared to individuals with high baseline perfusion, though the difference between groups did not meet our a priori level of significance (p=0.093). Individuals with low baseline perfusion also demonstrate a more rapid and hyperemic increase over the acute course of post-exercise recovery, exceeding baseline levels at 6-12, 12-18, and 18-24 minutes post-exercise (p<0.03). In contrast, hippocampal CBF did not exceed baseline levels at any time point in individuals with high baseline perfusion (p>0.10).

**Figure 3.**
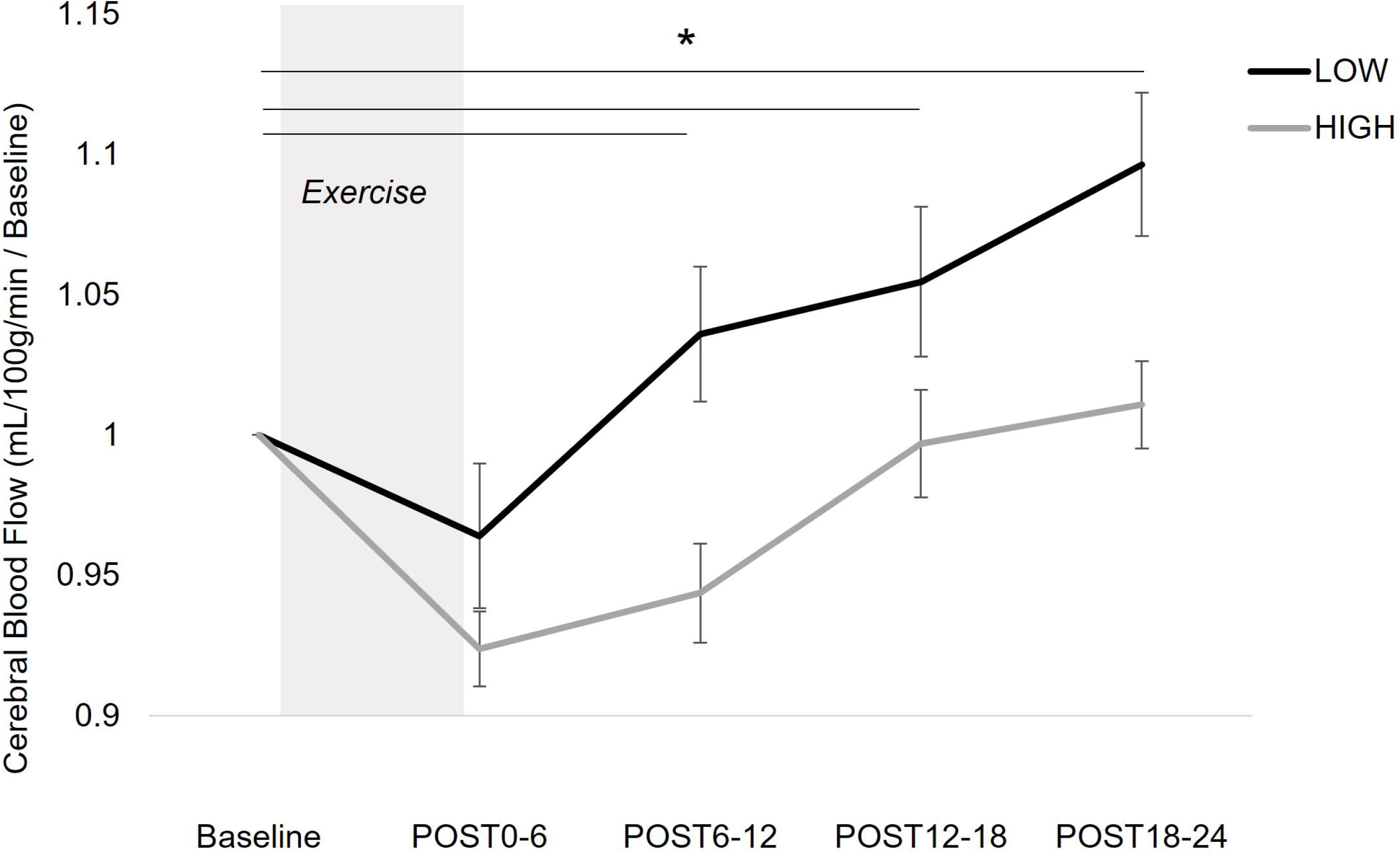
Hippocampal blood flow response to a bout of moderate-intensity aerobic exercise in older adults with low (< 26.00mL/100g tissue/min) and high (≥26.00mL/100g tissue/min) baseline hippocampal perfusion levels. Older adults with low baseline perfusion showed an attenuated immediate drop in hippocampal blood flow at 0-6 minutes post-exercise (POST0-6) followed by a more rapid and robust hyperemic rebound effect over the 24-minute assessment time course following exercise cessation, exceeding baseline levels at 6-12 (POST6-12) and 18-24 minutes (POST18-24) post-exercise (*p<0.03). Older adults with high baseline perfusion showed no increase in hippocampal blood flow above baseline levels at any timepoint assessed.

### 3.3. Acute exercise effects on cognitive performance and associations with hippocampal cerebral blood flow

Cognitive processing speed showed a main effect of time, increasing from baseline to post-exercise (X^2^ = 52.8, p<0.001), while memory performance showed no change (X^2^ = 2.8, p=0.10). There were no relationships between change in region-specific CBF and change in cognitive performance across either processing speed or memory domain tested (p>0.10).

## DISCUSSION

This study provides novel mechanistic insight into the acute global and region-specific cerebrovascular response to a bout of aerobic exercise in the aging brain. Our results yield three key findings: **(1)** brain regions showed a differential acute response to aerobic exercise, with the hippocampus *displaying a hyperemic rebound response* for at least 24-minutes post-exercise recovery; **(2)** older individuals with the *lowest regional perfusion* at baseline showed the *most rapid and robust increase in hippocampal blood flow* acutely following exercise cessation; and **(3)** acute exercise-induced changes in regional CBF were not associated with domain-specific increases in cognitive performance following exercise, suggesting dissociable cerebral mechanisms underpinning acute versus chronic improvements in cognitive function with exercise in older individuals. The present findings serve as a first step towards an improved understanding of the acute cerebrovascular response to aerobic exercise in the aging brain, laying a foundation for the development of precision-medicine techniques that target high-risk, older individuals with poor cerebrovascular health with aerobic exercise in the preclinical stages of disease.

### 4.1. Hippocampal blood flow rapidly and preferentially increases after exercise cessation in older adults

Following exercise cessation, hippocampal CBF increased more rapidly and robustly compared to other brain regions to exceed a post-exercise CBF above baseline at the end of the 24-minute post-exercise recovery assessment period (**Figure 2**). This acute region-specific cerebrovascular effect in older individuals is similar that reported in younger individuals (Steventon et al., 2020), who demonstrated region-specific increases in hippocampal blood flow acutely following a 20-minute bout of moderate-intensity aerobic exercise. The regional specificity of exercise-induced increases in hippocampal blood flow may be explained by local transient increases in norepinephrine and dopamine that occur following exercise (Dunn et al., 1996; Goekint et al., 2012; Meeusen & De Meirleir, 1995; Skriver et al., 2014) that have been demonstrated to trigger immediate local CBF modulation (Krimer et al., 1998). Repeated exposure to aerobic exercise stimuli over time can elicit cerebral angiogenesis (Ding et al., 2006) and increased hippocampal baseline pefursion (Pereira et al., 2007). In older individuals in the present study, acute hippocampal hyperemia following exercise is also consistent with findings of long-term exercise training, where hippocampal brain health and function show the greatest benefit compared to other brain regions (Erickson et al., 2011; Maass et al., 2015; Pereira et al., 2007; Seoane et al., 2022; van Praag et al., 2005). Here, the preferential hippocampal response to exercise may identify a cerebrovascular mechanism by which long-term exercise training elicits domain-specific cognitive benefits for memory function with aging (Ahlskog et al., 2011; Cotman & Berchtold, 2002; Jonasson et al., 2016; Kamijo et al., 2009; Maass et al., 2015; Makizako et al., 2015; Pereira et al., 2007; Vecchio et al., 2018; Zhang et al., 2020). For the first time, the present findings demonstrate the acute time scale of regional hippocampal blood flow responses to aerobic exercise in the aging brain, and implicate aerobic exercise as an effective behavioral approach to target early, age-related declines in hippocampal blood flow.

### 4.2. Poor cerebrovascular health elicits more rapid and robust increases in hippocampal blood flow acutely after exercise

When investigating the effects of regional cerebrovascular health on acute CBF response to exercise, we found that exercise-induced increases in hippocampal blood flow occurred only in older individuals with poor baseline perfusion, implicating an increased capacity for acute, exercise-induced vascular plasticity in the preclinical stages of age-related brain pathology. Lower baseline cerebral perfusion is an indicator of poor cerebrovascular health and associated with age-related pathology and disease (Iadecola, 2004; Ouellette & Lacoste, 2021; Weijs et al., 2022; Wolters et al., 2017). In older adults, we found the region-specific hippocampal blood flow effect after exercise (**Figure 2**) was driven by individuals with low regional perfusion; these individuals showed an attenuated immediate drop followed by a more robust, hyperemic rebound effect that quickly surpassed baseline levels and continued to increase over the acute 24-minute time course of exercise recovery (**Figure 3**). This effect of baseline regional perfusion level on acute, exercise-induced hippocampal blood flow response may be unique to aging, as there are no reports of this effect in similar investigations involving younger individuals (MacIntosh et al., 2014; Smith et al., 2010; Steventon et al., 2020). The higher prevalence of poor vascular health in older individuals as a result of aging processes (Alwatban et al., 2021; Thomas et al., 2013; Xing et al., 2017) may explain the specific aging context of this effect. This cerebrovascular health effect on acute CBF response to exercise is consistent with previous investigations of exercise training effectiveness as a function of cardiovascular health and fitness, where sedentary individuals with low fitness levels show the greatest health benefits in response to chronic exercise training (Ekelund et al., 2020; Ngandu et al., 2015). Here, our findings extend the principle that *individuals with poor vascular health possess the greatest potential for therapeutic exercise benefit* to the context of brain health with aging. Notably, the interactive effect between baseline perfusion and hippocampal blood flow response to exercise was present even after controlling for a proxy measure of fitness (i.e. exercise workload), suggesting that baseline regional perfusion may provide unique information for the prediction of exercise-induced vascular plasticity potential that is not gleaned from an individual’s level of physical fitness alone. Interestingly, post-exercise hippocampal blood flow never increased above baseline levels in individuals with high perfusion (**Figure 3**). As we only tested a single type of exercise, it is possible that a different type of exercise stimulus, for example high intensity interval training, may elicit more robust vascular and neural plasticity in these individuals (Boyne et al., 2019; Kaiser et al., 2022; Neva et al., 2022; Weston et al., 2022). The present results provide preliminary evidence for the use of baseline cerebrovascular perfusion as a predictive biomarker for brain health benefit from exercise in aging populations. Further, these findings implicate a sensitive time window in the early, preclinical stages of age-related disease processes for which exercise training may be most effective in high-risk older individuals.

### 4.3. Aerobic exercise elicits an acute drop in cerebral blood flow in older adults

A global and robust drop in CBF immediately following exercise cessation in older adults in the present study may reflect attenuated mechanisms of cerebrovascular regulation acutely post-exercise in the aging brain. We observed a sharp and immediate decrease in CBF across all brain regions from baseline levels during the 0-6 minute post-exercise time point (**Figure 2**). This CBF decline was likely triggered, at least in part, by the positional transition to supine at the start of the MR scan and may reflect cerebrovascular autoregulation mechanisms required during positional transfers (Claassen et al., 2021). Notably, the effect persisted even after controlling for mean arterial pressure, implicating the strong influence of cerebral-specific regulation mechanisms (Brassard et al., 2021). The global drop in CBF appears to be unique to older adult populations in the present study, as it was not reported younger individuals acutely after exercise, despite similar positional transfers required for MR acquisition (Smith et al., 2010; Steventon et al., 2020). The magnitude of CBF drop was also region-specific, showing an attenuated effect in the hippocampus (e.g. 6 % reduction in the hippocampus vs. 15% reduction in the superior parietal cortex) (**Figure 2**), particularly in individuals with low baseline perfusion (**Figure 3**). Interactions of aging and age-related vascular health with cerebrovascular autoregulation under behavioral conditions, e.g. aerobic exercise (Palmer et al., 2022) and positional transfers (Sorond et al., 2010; Whitaker et al., 2022), may explain the presence of this CBF hypoperfusion effect immediately after exercise in older adults in the present study. Older individuals may possess poor age-related cerebrovascular regulation under states of physiologic stress (Claassen et al., 2021; Ward et al., 2018), which are differentially regulated across brain regions and between individuals (Brassard et al., 2021). Acute exercise-induced CBF hypotension in older individuals warrants further investigation for interpretation and potential health implications in aging populations.

### 4.4. Acute cognitive performance and cerebral blood flow changes after exercise are not associated

In the present study, there was a lack of association between changes in cognitive performance and cerebral blood flow acutely post-exercise, suggesting dissociable acute effects of exercise on neural and vascular plasticity and elucidating the longer time scales necessary for the established link between neural and vascular plasticity in the cerebrum and hippocampus. Given the evidence for low regional CBF as a prognostic biomarker for the development of cognitive impairment and dementia (see Weijs et al., 2022 for comprehensive review and meta-analysis; Wolters et al., 2017) combined with evidence of aerobic exercise acutely benefiting cognitive function (Statton et al., 2015), it was surprising that we found no associations between change in domain-specific cognitive performance and region-specific CBF acutely after exercise. Together these findings suggest that repeated and chronic exposure to aerobic exercise training are needed to elicit vascular plasticity processes that positively influence cognitive function. The chronic time scales needed for exercise-induced cerebrovascular benefit on cognition may be consistent with other exercise-induced mechanisms involving the release of neurotrophic factors over longer time scales (e.g. weeks) for therapeutic benefit (Maass et al., 2015; Voss et al., 2013; Whiteman et al., 2014). It is possible that acute improvements in cognitive processing speed and memory were primarily due to task-specific practice effects, primarily driven by neural mechanisms (see Hubbard et al., 2009; Kleim & Jones, 2008 for review); in this case, exercise-induced vascular plasticity may play less of a role in this acute context. Future studies implementing a control condition without exercise will help dissociate practice from exercise-induced effects on increased cognitive performance.

### 4.5. Strengths and Limitations

Building upon prior human investigations in younger individuals and long-term exercise training effects on aging brain health, this work provides novel insight into *acute cerebral blood flow responses* to aerobic exercise in the *aging cerebrovascular system.* Our approach using arterial spin labeling MRI allowed investigation of magnetically labeled blood water molecules in the brain tissue capillary bed, enabling *region-specific* investigation of the acute time course of CBF response following aerobic exercise cessation.

Our focused investigation on the novel characterization of acute CBF responses with aging did not include a non-exercise control condition and should be considered carefully in the interpretation of these findings. The post-exercise MRI time window of our approach was based on acute hippocampal blood flow responses in younger individuals, who show hippocampal blood flow peaks within ~20 minutes after exercise (Steventon et al., 2020) and further enabled us to perform cognitive testing within <30 minutes post-exercise. However, this relatively shortened post-exercise time frame would not have captured vascular plasticity occurring on longer time scales post-exercise, warranting future research to extend the post-exercise MRI scan period to test for delayed blood flow effects in aging populations.

To date, a clinically-meaningful CBF threshold indicative of poor cerebrovascular health has not been established, as CBF has been assessed using different methods (e.g. single-photon emission computed tomography (SPECT), transcranial Doppler ultrasound (TCD), ASL-MRI)) (Alwatban et al., 2020, 2021; Kaufman et al., 2021; Vidoni et al., 2022; Ward et al., 2018; Weijs et al., 2022; Wolters et al., 2017). As a first step, the present study used the cohort median value for regional perfusion level to classify older individuals with “low” and “high” baseline perfusion; however, future studies are needed to empirically test the clinical significance of this baseline perfusion threshold as a metric of cerebrovascular health in larger cohorts of older adults and utilizing a standardized CBF assessment approach.

### 4.5. Conclusions

The present results reveal that older individuals can achieve region-specific hippocampal blood flow increases acutely following exercise similarly to those previously reported in younger individuals. We further identify a subgroup of older adults with low cerebrovascular health who show the most rapid and robust increase in hippocampal blood flow following exercise, suggesting older individuals in the preclinical stages of disease processes may have an increased capacity for exercise-induced regional vascular plasticity. Together, the present findings support an individualized, precision-based framework for the use of exercise to target brain health that carefully considers age-related changes in the cerebrovascular system.

## Acknowledgements

This work was supported by the National Institutes of Health R21 AG061548, P30 AG072973 and P30 AG035982, and the Leo and Anne Albert Charitable Trust. The Hoglund Biomedical Imaging Center is supported by a generous gift from Forrest and Sally Hoglund and funding from the National Institutes of Health including S10 RR29577, and UL1 TR002366.

## CONFLICTS OF INTEREST

The authors have no conflicts of interest with respect to bias of the research, authorship, and/or publication of this article.

## Notes

### Competing Interest Statement

The authors have declared no competing interest.

